# Sampling design and sample processing affect soil biodiversity assessments

**DOI:** 10.64898/2026.01.05.697626

**Authors:** Meirong Chen, Olesya Dulya, Vladimir Mikryukov, Ovidiu Copot, Martin Metsoja, Leho Tedersoo

## Abstract

Biodiversity surveys require an appropriate sampling design for optimal performance and comparability across space and time and across studies. Based on PacBio and Illumina amplicon sequencing of animals, bacteria and fungi, we assessed and compared various soil sampling designs from widely used continental and global metabarcoding-based biodiversity projects. Sampling designs revealed up to 27-fold, 6-fold and 15-fold differences in biodiversity estimates for animals, bacteria and fungi, respectively. Taxonomic coverage depended mostly on the number of subsamples but not sampling area within the 347–1790 m^2^, 428–1924 m^2^, 327–1790 m^2^ plot size range for animals, bacteria and fungi, respectively. Additional sampling of subsoil did not add significantly to diversity estimates of animals and fungi, although there was a slight positive effect on bacteria. Both soil pooling (compositing subsamples before DNA extraction) and DNA pooling (combining DNA extracts prior to PCR) reduced differences between sampling designs by decreasing diversity estimates in designs with many subsamples while increasing them in designs with fewer subsamples. However, pooling did not eliminate the influence of sampling factors (soil depth, sampling area and sample size). DNA pooling outperformed soil pooling in inventorying animals and fungi but not bacteria. Pooling had no effect on recovering rare biological species or sequencing artefacts. Soil pooling saved from 79.5% to 98.3% of labour and analytical costs compared with no pooling, depending on the number of pooled subsamples. The remarkable impact of sampling design on community diversity and composition should be considered during data collection and meta-analyses compiling data from different sampling designs.

## Introduction

Determining a proper sampling design is a fundamental issue in biodiversity surveys, which relates to research questions, target organisms and resources (Bruckner et al., 2000; Querner & Bruckner, 2010; Ranjard et al., 2003). Suboptimal sampling designs may bias biodiversity estimates, depending on the spatial pattern of sampling, sample size, sample volume and sampling area (Bell et al., 2005; Chase & Knight, 2013; Kang & Mills, 2006; MacArthur, 1965; Rahbek, 2005; Tan et al., 2017). Furthermore, 95% of sampling strategies are non-reproducible due to subjectivity or insufficient description of the site selection or sampling process (Dickie et al., 2018; Whitcomb & Stutz, 2007).

The largely stochastic community assembly, spatial aggregation and enormous taxonomic richness make the selection of sampling designs for soil microbial diversity research more complicated than for macroorganisms (Fukami, 2010; Green, 1979; Kang & Mills, 2006; Lemoine et al., 2023; Lücking et al., 2021). Appropriate sampling is particularly relevant in soil metabarcoding surveys due to the patchy distribution of species and our capacity to sample only a tiny fraction of the community. Research teams have reached different trade-offs between sampling comprehensiveness and use of time in the field and lab and financial resources (Bach et al., 2018; Engel et al., 2012; Fierer & Jackson, 2006; Guerra et al., 2021), but these decisions are typically unguided by the real data or power analyses.

To reduce the laboratory costs while maintaining representativeness, sampling designs involving subsampling are common in molecular ecology. The subsamples are usually pooled before or after DNA extraction to reduce the costs of sample processing and molecular analyses (Dickie et al., 2018). However, sample pooling has raised several technical concerns about equal subsample quality and quantity and overall dilution. Negative impacts of soil and DNA sample pooling on diversity estimates have been reported in ARISA (Automated ribosomal intergenic spacer Analysis) and T-RFLP (Terminal restriction fragment length polymorphism) for bacteria, fungi and animals (Engel et al., 2012; Manter et al., 2010; Osborne et al., 2011). Studies based on high-throughput sequencing (HTS) technologies that provide a higher resolution in species identification also show that pooling effect varies among taxonomic groups, habitats, and diversity measures. In particular, soil pooling has caused loss of fungal species richness estimates but not on Shannon diversity index (Song et al., 2015). Pooled samples enabled detection of fewer arthropods from soil (Porter et al., 2019). It is crucial to understand how different sampling design features – such as sampled area, sample number, depth and subsample pooling – affect diversity estimates of soil organisms. This would provide a source for information-driven decisions about an optimal sampling design considering sampling representativeness, time and budgetary constraints.

In this study, we assessed the effects of sampling design and pooling strategies in recovering the richness of soil-inhabiting bacteria, fungi and animals based on PacBio and Illumina amplicon sequencing. The study had three principal aims: 1) to evaluate the relative performance of existing, broadly used sampling designs in recovering biodiversity of different organism groups; 2) to estimate the effect of pooling either soil before DNA extraction or DNA extracts before the PCR step on species recovery; and 3) to provide information for decision making when considering sampling design and analytical strategies related to pooling. We expected that sampling designs with more subsamples covering a larger area would reveal more species (Green et al., 2004), especially in soil animals, that typically have patchy distribution (Berg, 2012). We also suggested that the effect of pooling either soil or DNA depends on sampling design. In particular, we expected a stronger species loss for designs with larger sample sizes due to the dilution effect (Manter et al., 2010; Morita et al., 2021). We also calculated the richness and Shannon diversity index at various sequencing depths to determine whether the resources saved by sample pooling could be reallocated to achieve greater sequencing depth, thereby increasing the probability of detecting low-abundance species.

## 2 Methods

### 2.1 Sampling designs and field work

We conducted soil microbiome surveys in three natural forest sites growing on luvisols in Estonia (Tartu county). The Järvselja site (58°16’N, 27°19’E) represents an old-growth (140 years) mixed forest dominated by *Tilia cordata* and *Picea abies*. The Kambja site (58°14’N, 26°42’E) is a *Betula pendula* and *Picea abies* dominated 120-year-old forest. The Vasula site (58°28’N, 26°44’E) is a dense *Quercus robur* and *Fraxinus excelsior* dominated woodland (140 years). In each forest site, we located a relatively homogeneous 10,000 m^2^ area and established a randomly selected point at least 3 m distant from the nearest tree (DBH > 10 cm) with a uniform microtopography within 1 m radius. Then we delimited the maximum sampling area (2500 m^2^) around this central core. These points represented plot centres for all compared sampling designs (Table 1, Fig. 1). The samples were collected following the original protocols (see the referenced works in Table 1), using either a 50-mm PVC pipe, a sharp knife or a teaspoon. In each site, soil sampling was performed in dry weather on a single day in July 2021. To partition the effects of the sampling area and sample size, we modified the N_40_D_0-5A_ (40 subsamples of 0–5 cm depth in 2500 m^2^) original design (Tedersoo et al., 2021) by increasing the number of subsamples to 62 (N_62_D_0-5_) or reducing the sampling area to 1400 m^2^ (N_40_D_0-5B_). For the N_9_D_0-10_ design used for Soil BON (Guerra et al., 2021), we collected four additional soil cores for the OTU richness estimates in different sampling depths from the same 900-m^2^ plot. For all N_9_D_0-10_ samples, we additionally collected a subsoil sample from 30–40 cm depth that was treated either separately (N_9_D_30-40_) or combined with the topsoil samples (N_9_D_0-10_). We recorded the coordinates of each individual sample or subplot by using tape measurements from a central spot and two crossing axes. In total, we collected 321 soil cores across the three sampling sites.

**Figure 1.**
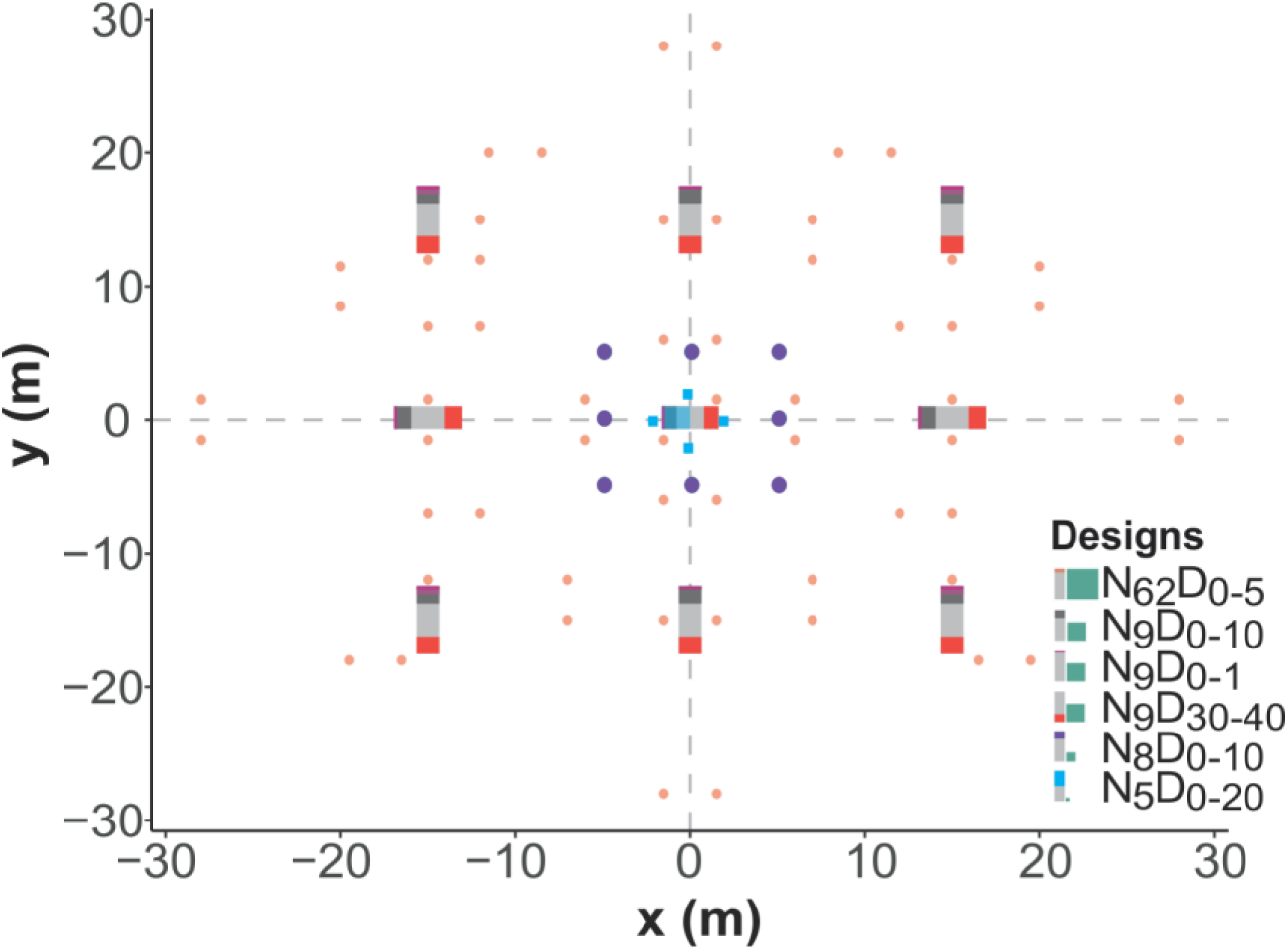
The scheme of sampling designs. Colours indicate different sampling designs. In legend symbols, the green rectangle is proportional to the sampling area and colour shadow in a gray bar denotes sampling depth. The columns in the map show the sampling depth of designs at overlapping core positions.

**Table 1.**
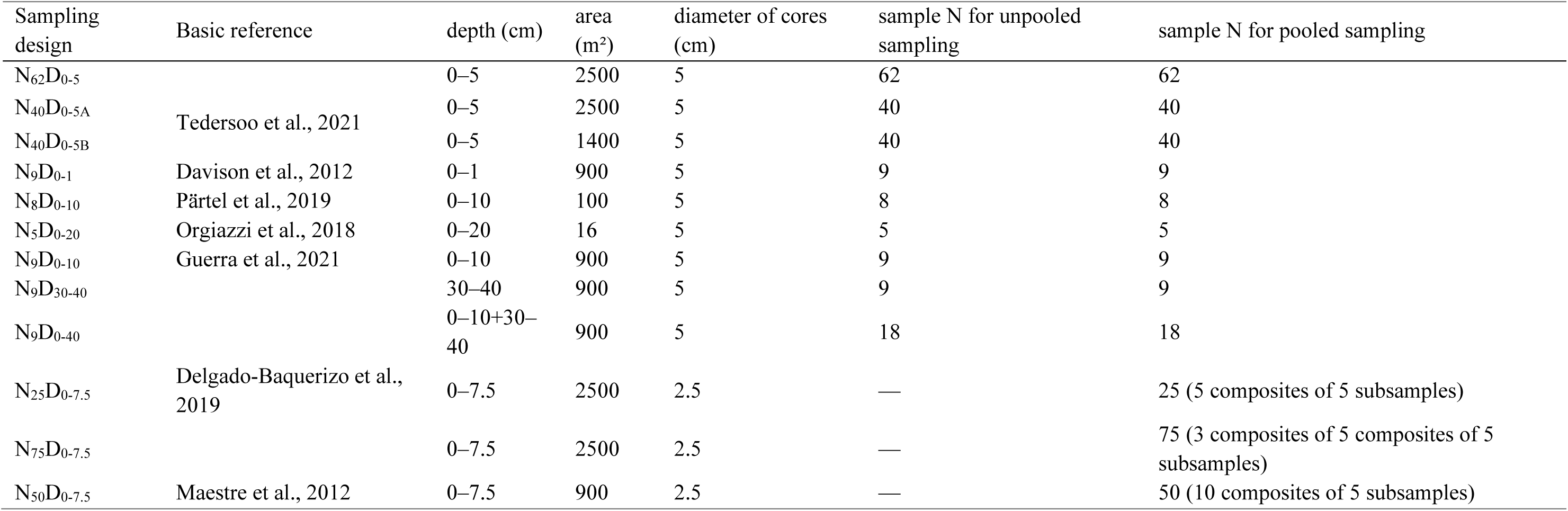
Types of sampling designs and pooling strategies involved in the experiment. In the code for sampling design, N represents the number of subsamples and D denotes the sampling depth range (cm).

| Sampling design | Basic reference | depth (cm) | area (m<sup>2</sup>) | diameter of cores (cm) | sample N for unpooled sampling | sample N for pooled sampling |
| --- | --- | --- | --- | --- | --- | --- |
| N<sub>62</sub>D<sub>0-5</sub> |  | 0–5 | 2500 | 5 | 62 | 62 |
| N<sub>40</sub>D<sub>0-5A</sub> | Tedersoo et al., 2021 | 0–5 | 2500 | 5 | 40 | 40 |
| N<sub>40</sub>D<sub>0-5B</sub> |  | 0–5 | 1400 | 5 | 40 | 40 |
| N<sub>9</sub>D<sub>0-1</sub> | Davison et al., 2012 | 0–1 | 900 | 5 | 9 | 9 |
| N<sub>8</sub>D<sub>0-10</sub> | Pärtel et al., 2019 | 0–10 | 100 | 5 | 8 | 8 |
| N<sub>5</sub>D<sub>0-20</sub> | Orgiazzi et al., 2018 | 0–20 | 16 | 5 | 5 | 5 |
| N<sub>9</sub>D<sub>0-10</sub> | Guerra et al., 2021 | 0–10 | 900 | 5 | 9 | 9 |
| N<sub>9</sub>D<sub>30-40</sub> |  | 30–40 | 900 | 5 | 9 | 9 |
| N<sub>9</sub>D<sub>0-40</sub> |  | 0–10+30–40 | 900 | 5 | 18 | 18 |
| N<sub>25</sub>D<sub>0-7.5</sub> | Delgado-Baquerizo et al., 2019 | 0–7.5 | 2500 | 2.5 | — | 25 (5 composites of 5 subsamples) |
| N<sub>75</sub>D<sub>0-7.5</sub> |  | 0–7.5 | 2500 | 2.5 | — | 75 (3 composites of 5 composites of 5 subsamples) |
| N<sub>50</sub>D<sub>0-7.5</sub> | Maestre et al., 2012 | 0–7.5 | 900 | 2.5 | — | 50 (10 composites of 5 subsamples) |

Each sample was placed into a sterile paper bag and dried within 12 hours of collection at 35 ℃ in a drying cabinet. The dried soil samples were initially crushed by rubbing the plastic bag, and ca. 1 g of the finest particles were further homogenised using tungsten beads in a Retsch MM400 homogeniser (Retsch GmbH, Haan, Germany) at 30 Hz for 5 min. Then, 0.2 g of homogenised soil was weighed for DNA extraction. DNA extraction for all samples was performed using the PowerMax Soil DNA Isolation kit (Qiagen, Carlsbad, CA, United States) following the manufacturer’s instructions. To minimise contamination, we wore disposable gloves and cleaned instruments and work surfaces with 70% ethanol throughout all sample-processing steps.

We focused on three sample pooling strategies for each site, including 1) unpooled strategy, 2) pooling of soil and 3) pooling of DNA. The unpooled strategy involves separate DNA extraction and analysis of individual samples as described above. For the DNA pooling strategy, 2 μL of each DNA extract from the samples collected within the same sampling design were mixed into a DNA pool. For the soil pooling strategy, ca. 3 g of the hand-crushed samples collected within the same design were taken to the composite sample, followed by mixing, homogenisation by bead beating and DNA extraction from 0.2 g of material.

### 2.3 PCR and sequencing

Soil samples were analysed for the animal, fungal and bacterial biodiversity based on DNA metabarcoding. We conducted PCR amplification using the universal eukaryotic primers ITS9mun (5’ GTACACACCGCCCGTCG 3’) and ITS4ngsUni (5’ CGCCTSCSCTTANTDATATGC 3’) for amplification of the fungal Internal Transcribed Spacer (ITS) marker following Tedersoo et al. (2021). The primers COI.intF (5’ GGWACWGGWTGAACWGTWTAYCCYCC 3’) and COI.2198 (5’ TAIACYTCIGGRTGICCRAARAAYCA 3’) were used for amplifying the mitochondrial Cytochrome Oxidase I (COI) gene following Anslan et al. (2021). The primers 515F (5’ GTGYCAGCMGCCGCGGTAA 3’) and 926R (5’ GGCCGYCAATTYMTTTRAGTTT 3’) were used to amplify the bacterial 16S rRNA gene as described in Bahram et al. (2018). The PCR products were visualised on a 1% agarose gel. DNA concentrations were assessed using Qubit 3.0 (Thermo Fisher Scientific, Chicago, USA).

The PCR products of the respective libraries were pooled based on the product strength on a gel, by mixing 0.5 μL to 20 μL of PCR product. The mixtures were shipped to a service provider for library preparation and sequencing. The animal COI and bacterial 16S regions were sequenced on an Illumina NovaSeq 6000 at the Novogene (Cambridge, United Kingdom), whereas the fungal ITS region was sequenced on a PacBio Sequel II at the Norwegian Sequencing Centre, University of Oslo (Oslo, Norway). Illumina NovaSeq 2x250 paired-end protocol was used for sequencing the COI and 16S amplicons. The fungal samples producing < 2000 reads and bacterial and animal amplicons with < 50,000 reads were subjected to resequencing in another sequencing library.

### 2.4 Bioinformatics

For each group (fungi, bacteria and animals), we followed current best practices and applied state-of-the-art analytical workflows. Bioinformatics analysis of sequence data of the ITS region was conducted using the NextITS pipeline v.0.5.0 (Mikryukov et al., 2025). The fungal sequences were demultiplexed with LIMA v.2.7.1 (Pacific Biosciences; https://lima.how/) using the minimum barcode score of 93. Full-length ITS sequences were extracted using ITSx v.1.1.3 (Bengtsson-Palme et al., 2013). Chimeric sequences were removed using a two-step approach: *de novo* detection using the UCHIME algorithm (Edgar et al., 2011) with a maximum chimera score of 0.6 (Nilsson et al., 2015) and a reference-based method using the EUKARYOME database v.1.7 (Tedersoo et al., 2024). Sequence clustering was performed using VSEARCH v.2.28.1 (Rognes et al., 2016) at a 98% sequence similarity threshold (Bradshaw et al., 2023). For post-clustering curation, we used the LULU algorithm (Frøslev et al., 2017) with a sequence similarity threshold of 95%. To taxonomically annotate fungal and bacterial sequences, we used the Naïve Bayes classifier (Wang et al., 2007) implemented in QIIME2 v.2023.9.1 (Bolyen et al., 2019) along with the EUKARYOME reference database. Sequences were assigned to the lowest taxonomic level for which the classification bootstrap confidence was at least 80% (Bokulich et al., 2018). Non-fungal OTUs and sequences unclassified at the kingdom level were excluded from further analyses.

The bacterial dataset was analysed using DADA2 v.2023.9.0 (Callahan et al., 2016) as implemented in the QIIME2 plugin. We removed the forward reads < 219 bp and reverse reads < 216 bp as low-quality sequences. The reads were clustered using a 97% sequence similarity threshold (Liu et al., 2023). We used a weighted taxonomic classifier based on the SILVA database v.138.1 (Kaehler et al., 2019; Quast et al., 2013; Robeson et al., 2021). Unclassified, non-target (e.g., mitochondrial and plastid) and eukaryotic OTUs were excluded from the dataset. OTUs with a taxonomic confidence value < 80% at the kingdom level were removed (Bokulich et al., 2018).

For the analysis of COI amplicons, ASVs were obtained using MetaWorks v.1.13.0 (Porter & Hajibabaei, 2022) with default settings. OTUs were calculated based on sequence clustering at a 99% sequence similarity threshold (Zhang & Bu, 2022). Taxonomic annotation of OTUs was conducted using the RDP Classifier v.5.1.0 (Wang et al., 2007). Only OTUs assigned to Rotifera, Nematoda, Arthropoda and Annelida were retained, and those with a phylum-level taxonomic confidence value below 80% were removed (Bokulich et al., 2018).

To understand the contribution of potential artefacts and biological species to the pooling effect, we focused on the three largest fungal groups, *Thelephoraceae*, *Sebacinaceae* and *Russulaceae*, where we had some taxonomic expertise. Representative sequences of each OTU and best-matching reference sequences were aligned using MAFFT v.7.525 (Katoh & Standley, 2013) with default options. The first step in artefact recognition was performed using Aliview v.1.30 (Larsson, 2014). The putative artefacts included sequences with i) long indels (>20 bases throughout the sequence); and ii) more than one single-nucleotide indel in the 200-base window spanning the entire 5.8S rRNA gene and first part of ITS2. The remaining sequences were subjected to Maximum Likelihood tree construction using IQ-Tree v.2.1.4 (Minh et al., 2020) using default options. In the second step of artefact recognition, taxa with relatively long branches compared with the best hit, corresponding to > 2% sequence divergence, were removed.

Finally, we removed the samples with <100 reads, retaining 317 individual subsamples for bacteria, 310 for fungi and 295 for animals. We retained all 36 DNA-pooled samples for animals, bacteria and fungi, as well as 36 soil-pooled samples for animals and bacteria and 35 soil-pooled samples for fungi.

### 2.5 Statistical analysis

All statistical analyses were conducted in R v.4.3.1 (R Core Team, 2023). Sample completeness was estimated using Good’s coverage (Chiu & Chao, 2016; Kortmann et al., 2025). Coverage was near-saturated (animals: 1.00 [median] ± 0.00 [SD], bacteria: 1.00 ± 0.0003; fungi: 0.96 ± 0.04). Therefore, to control for sequencing effort, analyses of the effects of sampling design, pooling strategy, or sampling intensity (number of samples) on community diversities were run on sequencing depth-standardised data. For animals, bacteria and fungi separately, OTU tables were rarefied to the near-minimum read number (animals: 880; bacteria: 1067; fungi: 814 in unpooled samples; animals: 15962; bacteria: 39290; fungi: 14409 in pooled samples). To reduce rarefaction stochasticity, we performed multiple rarefactions (1000 iterations) and averaged the resulting diversity estimates. This analysis was performed using metagMisc v.0.5.0 (Mikryukov, 2025) and phyloseq packages v.1.44.0 (McMurdie & Holmes, 2013).

#### 2.5.1 Effect of sampling design and pooling strategy on diversity

Rarefied OTU accumulation curves were constructed using the vegan package v.2.6-4 (Oksanen et al., 2015). Cumulative abundance curves were built to show the OTU detection ability of each design. We compared the mean (within-sample) OTU richness across the unpooled sampling designs using a mixed linear model, with the site as a random factor. For pooled sampling designs, the site was treated as a fixed factor. We compared the soil and DNA pooling in diversity estimates based on the mixed linear model where site was included as a random effect and sampling design as a fixed effect. These analyses were performed using the marginaleffects package v.0.18.0 (Arel-Bundock V, 2024) based on the model built by the lme4 package v.1.1-35.1 (Bates et al., 2015).

To test the effect of unpooled sampling design, we compared the obtained total richness values (within-site) of each unpooled sampling design based on a linear model with the sampling sites and sampling designs as fixed factors using the marginaleffects and lme4 packages. We estimated sample coverage and extrapolated site-richness using species-by-sampling-site-unit matrices as inputs based on the iNEXT.3D v.1.0.4 package (Chao et al., 2021). We regressed species richness on the sampling area using a linear mixed model, with the site as a random factor, implemented in the lmerTest v.3.1-3 package (Kuznetsova et al., 2017). Eight samples (4 pairs) were selected from the N_62_D_0-5_ sampling design to test the sampling area effect on richness.

We performed the permutational multivariate analysis of variance (PERMANOVA; Anderson, 2017) to compare community composition across sampling designs based on the Bray-Curtis distances (vegan package) among individual samples. To assess the effect of sampling design on beta-diversity estimates, we conducted the homogeneity test of multivariate dispersion at each site using the Bray-Curtis distance using the vegan package. To test the impact of the sample pooling on community composition similarity among sampling designs, we performed the marginal contrast with site as a random factor and designs as a fixed factor based on Bray-Curtis distance using vegan, marginaleffects, and lme4 packages.

#### 2.5.3 Pooling effect analysis

We define the pooling effect as the ratio of OTU richness or effective number of OTUs (Jost, 2006) between pooled and unpooled sampling obtained with the same sampling design. Although the Shannon index is widely used, its logarithmic basis makes interpreting its ratios unintuitive. Exponentiating the Shannon index yields an effective number of OTUs, corresponding to Hill number of q = 1 and representing the number of equally abundant species required to achieve the observed diversity and facilitating more intuitive comparisons among sampling designs. To compare the pooling effects among various sampling designs, the summary sequencing depth of each sampling design (the total reads of the same unpooled design) was used to assess the pooling effect within each design (Table S1). We compared the effects of pooling strategy (soil vs. DNA) using a linear mixed-effects model, with sampling site and sequencing depth as random effects, sampling design and pooling strategy as fixed effects. The analyses were conducted with the lme4 and marginaleffects packages. The effect sizes of the fixed effects were estimated as the partial eta-squared (η_p_²) using the effectsize v.0.8.6 package (Ben-Shachar et al., 2020).

To assess the artefacts and rare species attribution to the pooling effect, we compared the proportion of artefact (N(artefacts) / N(OTUs)), rare OTUs (N(rare) / N(OTUs)) and rare artefact (N(artefacts) / N(rare species)) and the number of unique OTUs, rare artefacts, and unique artefacts in *Thelephoraceae*, *Sebacinaceae* and *Russulaceae*. The comparison was conducted using a mixed linear model with sampling designs, and the pooling strategy as fixed factors and the sampling site as a random factor based on the lme4 and the marginaleffects package. Model parameters and performance were assessed using the parameters v.0.21.6 and performance v.0.11.0 packages <u>(Lüdecke et al., 2020, 2021)</u>. Species with relative abundance ≤ 0.05% of total reads were defined as rare species.

We evaluated the extent of resource savings achieved through the pooling strategies using calculator available at https://mycology-microbiology-center.github.io/Soil_biodiversity_assessments/. In this work, the pooling effect for DNA and soil pools was evaluated at varying sequencing depths, reflecting the proportions of the expenses for sequencing a pooled sample relative to the sequencing of all unpooled samples within the same design. This analysis highlights how pooling strategies can reduce resource consumption while maintaining effective community composition and diversity assessments. The costs are measured in working hours and units of chemicals used for DNA extraction, PCR and sequencing. One working hour corresponds to roughly 10 extracted DNA samples or 40 PCR reactions (including product check and purification). We used a mixed linear model to analyse the pooling strategy’s impact on the pooling effect, where the site, sequencing depth and sampling designs were treated as random factors using the lme4 and marginaleffects packages.

## 3 Results

### 3.1 Individual sample diversity varies across sampling designs

Diversity estimates in individual soil samples differed across sampling designs, depending on sampling depth and target taxon (Fig. 2A; Table S2). The highest animal OTU richness was obtained with the N_9_D_0-1_ and N_62_D_0-5_ designs, with a gradual decrease through the N_8_D_0-10_ and N_9_D_0-10_ design to N_5_D_0-20_ and N_9_D_30-40_ designs. Bacterial OTU richness was significantly higher in the N_62_D_0-5_ and N_40_D_0-5A_ designs compared with all the other designs. The bacterial Shannon diversity index gradually decreased from the designs sampling within 0-5 cm through the designs sampling within 0-10 cm to N_5_D_0-20_ and N_9_D_30-40_. Conversely, fungal OTU richness did not significantly differ among sampling designs, except for much lower values in the N_9_D_30-40_ design.

**Figure 2.**
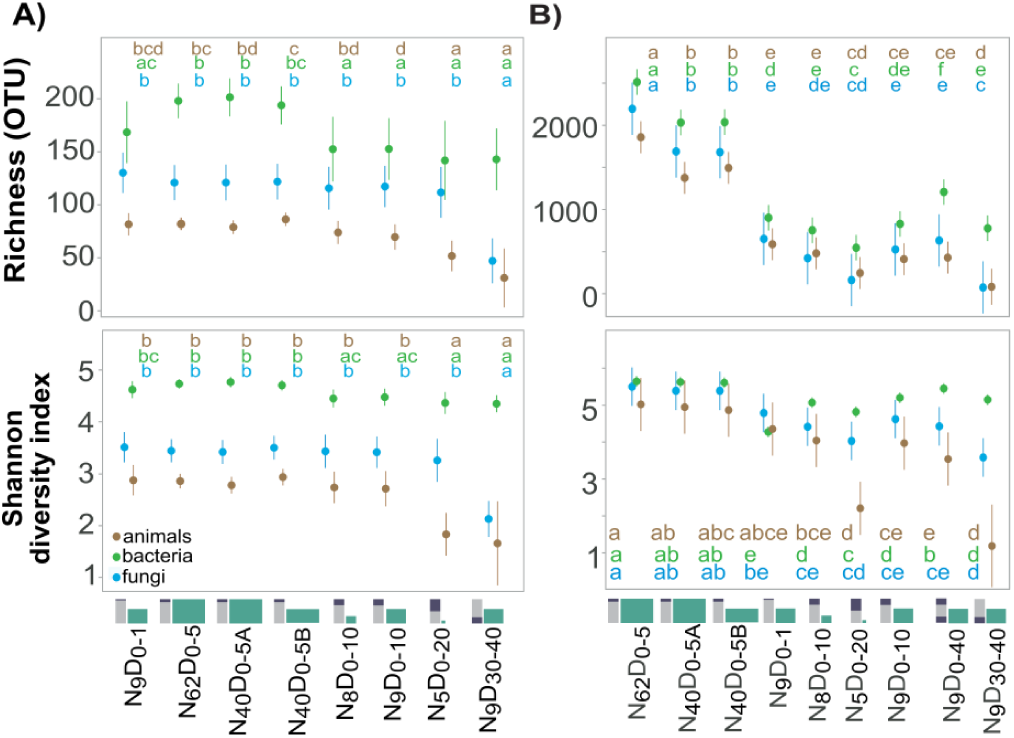
Diversity of soil organisms across sampling designs in individual samples. A) Marginal estimates of mean OTU richness and the Shannon diversity index in individual samples across sampling designs. (B) Marginal estimates of mean within-site OTU richness and Shannon diversity index. Similar letters group sampling designs with mean values that do not differ significantly (α = 0.05).

Based on PERMANOVA tests, community composition dissimilarity among sampling designs was slightly but significantly greater than within the designs (animals: F_(7, 449)_ = 1.3, R^2^ = 0.02, p = 0.002; bacteria: F_(7, 531)_ = 3.4, R^2^ = 0.04, p = 0.001; fungi: F_(7, 459)_ = 1.5, R^2^ = 0.02, p = 0.001; Tables S3, S4). Animal communities of the N_5_D_0-20_ samples significantly differed from those collected from the uppermost topsoil (N_9_D_0-1_ and N_62_D_0-5_ designs); however, no significant differences were observed aside from these. Bacterial communities of 0–1 cm, 0–5 cm, 0–20 cm and 30–40 cm depth all differed from each other but (except for N_9_D_30-40_ and N_9_D_0-1_) overlapped with the designs assessing 0–10 cm depth. In fungi, samples obtained from 0–1 cm, 0–5 cm, 0–10 cm and 0–20 cm depth had generally similar fungal communities, except for significant differences in the 0–1 cm topsoil compared with 0–20 cm. Subsoil fungal communities of N_9_D_30-40_ samples differed from other sampling designs, except the N_5_D_0-20_ design (Table S3).

Sampling designs significantly differed in the variability of community composition (Table S4), with the lowest dispersion in N_5_D_0-20_ (animals: 0.236, bacteria: 0.339, fungi: 0.326). The N_62_D_0-5_ (0.670), N_9_D_30-40_ (0.447) and N_62_D_0-5_ (0.597) showed the highest dispersion in the composition of animals, bacteria and fungi, respectively.

### 3.2 Sampling design influences overall biodiversity estimates

The total OTU richness in the site, assessed based on unpooled sampling, was the highest for the topsoil sampling designs (depth: 0–5) with a high number of subsamples (Fig. 2B; see Table S2). The richness ratios between sampling designs reached 26.8, 5.9 and 14.7, for animals, bacteria and fungi, respectively. Specifically, the N_40_D_0-5_ design yielded, respectively, 2.4–22.1, 1.7–4.8 and 1.7–11.5 times more animal, bacterial and fungal OTUs than designs with fewer subsamples (subsoil N_9_D_30-40_ was excluded from ratio calculation). The N_5_D_0-20_ and N_9_D_30-40_ designs generally revealed significantly lower numbers of OTUs per site compared to the other designs.

Compared to the N_9_D_0-10_ design, the N_9_D_0-40_ design (i.e., merging topsoil and subsoil samples) added on average 4.8%, 54.1% and 17.3% to OTU richness of animals, bacteria and fungi, respectively; yet these values were significantly below the values of N_62_D_0-5_ and N_9_D_0-1_ designs, suggesting that an extra effort of sampling subsoil or deep cores does not add much to the recovered richness, except in bacteria. Across all sites, subsoils harboured only 0.6%, 4.9% and 1.5% of unique OTUs of animals, bacteria and fungi, respectively.

In all organism groups, the Shannon diversity index varied similarly to richness, decreasing from the topsoil samples (depth: 0–5 cm) through the N_9_D_0-1_, N_9_D_0-10_ and N_8_D_0-10_ designs to the N_5_D_0-20_ and N_9_D_30-40_ designs (Fig. 2B; see Table S2). The ratios of effective numbers of OTUs between N_40_D_0-5_ and other designs with fewer subsamples (excluding N_9_D_30-40_) were 1.1–60 for animals, 1.1–4.7 for bacteria and 1–6.9 for fungi. For all groups of organisms, the topsoil samples detected more low-abundance OTUs and a lower proportion of dominant OTUs than those from the other designs (Figs. S1, S2). The N_5_D_0-20_ and N_9_D_30-40_ unpooled sampling designs captured the fewest low-abundance OTUs.

We further analysed if sampling design affects the patterns of diversity distribution across sampling sites (Fig. 3A). For animal richness, site ranking performed with N_5_D_0-20_ and N_8_D_0-10_ deviated from the other designs (Fig. 3A). For fungi, the designs involving 0–5 cm depth ranked the sites in the order Kambja-Vasula-Järvselja, while those sampling within 0–10 and 0–20 cm, ranked in the order Kambja-Järvselja-Vasula. Designs sampling from the 0–1 cm and 30–40 cm depth deviated from both above patterns in fungi. In bacterial richness estimates, all sampling designs, except N_8_D_0-10_, ranked sites identically.

**Figure 3.**
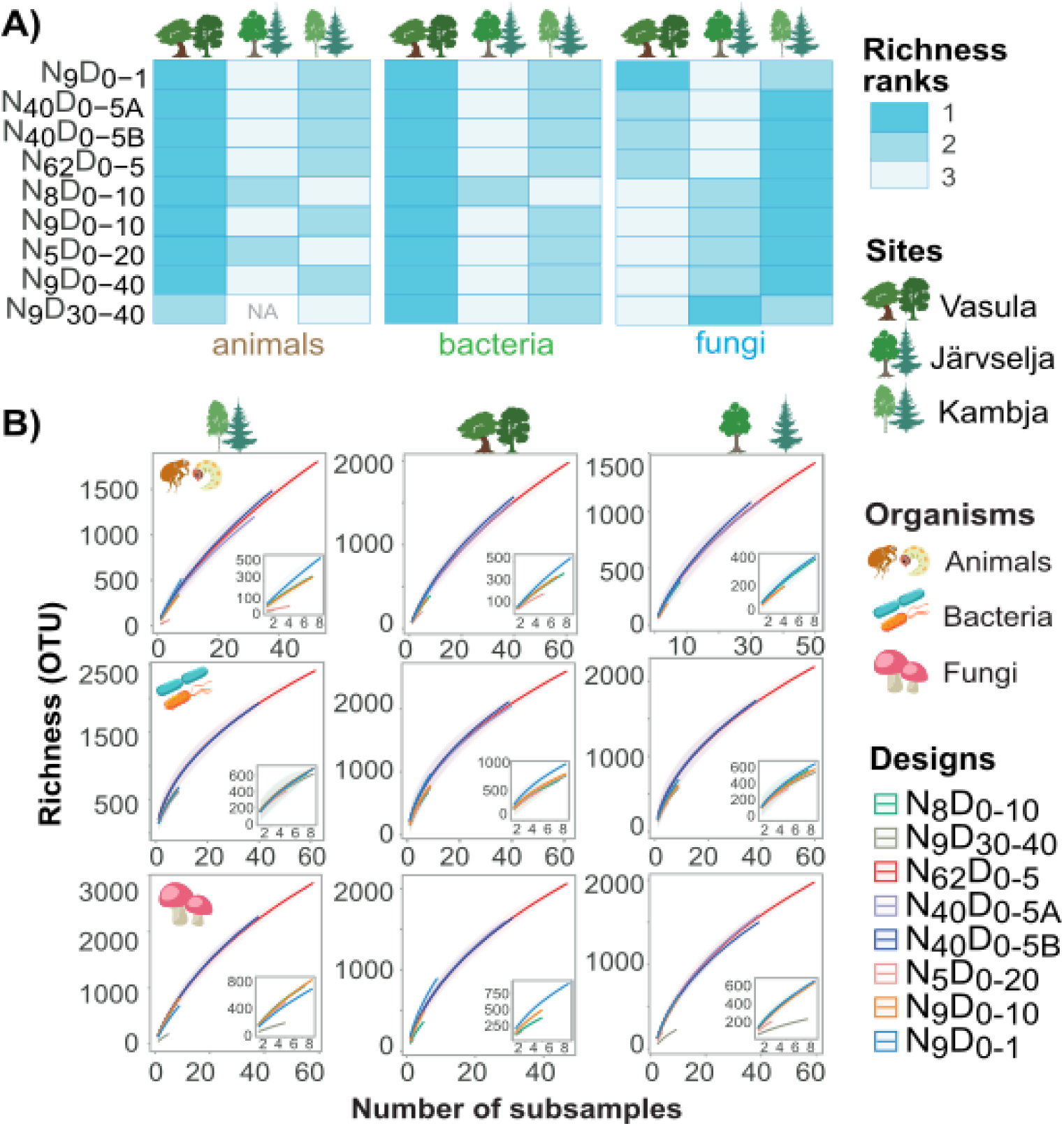
Relative performance of sampling designs in analyzing within-site diversity. A) Within-site richness ranked within each sampling design. B) OTU accumulation curves with sampling effort for different sampling designs.

For most microbial groups, the rate of OTU accumulation with sampling effort was the greatest for the designs focusing on the upper soil layers (Fig. 3B). Sampling only the subsoil (N_9_D_30-40_ design) yielded the lowest rate of OTU gain for animals and fungi, while bacterial richness accumulation was slowest in the N_5_D_0-20_ design.

We specifically tested the importance of sampling area on richness recovery by subsampling 8-sample combinations (N(animal) = 35; N(bacteria) = 47; N(fungi) = 31) within the N_62_D_0-5_ design while varying the rectangular (|width – length| ≤ 5 m) sampling area from 327 m^2^ to 2000 m^2^. The sampling area had no effect on OTU richness in any of the organism groups (p > 0.05 in all cases; Table S5). Based on all topsoil samples, animals, bacteria and fungi exhibited a significant positive spatial autocorrelation in community composition (Fig. S3).

### 3.3 Sample pooling reduces differences among sampling designs

To compare the sampling designs in diversity estimates obtained with pooled samples, we rarefied the samples at near-minimal sequencing depths, i.e. at 15,962, 39,290 and 14,409 sequences for animals, bacteria and fungi, respectively. Similarly to unpooled samples, pooled samples of the N_62_D_0-5_ design revealed the highest numbers of OTUs and the highest Shannon index (Fig. 4A, B; Table S6) due to a greater number of low-abundance OTUs (see Figs. S1, S2). However, the variability of richness and Shannon diversity index across pooling sampling designs was lower than across sampling designs with unpooled sampling. Specifically, in pooled samples, the richness ratio between N_40_D_0-5_ and other designs (excluding N_9_D_30-40_) were 0.9–3.1, 0.8–2.2 and 0.9–3.5 folds for animals, bacteria and fungi, respectively. The ratios of effective numbers of OTUs were 0.3–20, 0.8–2.4 and 0.7–5.8 folds for animals, bacteria and fungi, respectively. Compositional similarity between sampling designs was higher with pooled sampling than with unpooled sampling for bacteria (marginal contrast MC_UNPOOLED-SOIL_ = −0.05, p = 0.047; MC_UNPOOLED-DNA_ = −0.06, p = 0.005; see Table S5) and fungi (MC_UNPOOLED-SOIL_ = −0.08, p < 0.01; MC_UNPOOLED-DNA_ = −0.11, p < 0.001) but not for animals (MC_UNPOOLED-SOIL_ = 0.04, p = 0.2; MC_UNPOOLED-DNA_ = −0.04, p = 0.2).

**Figure 4.**
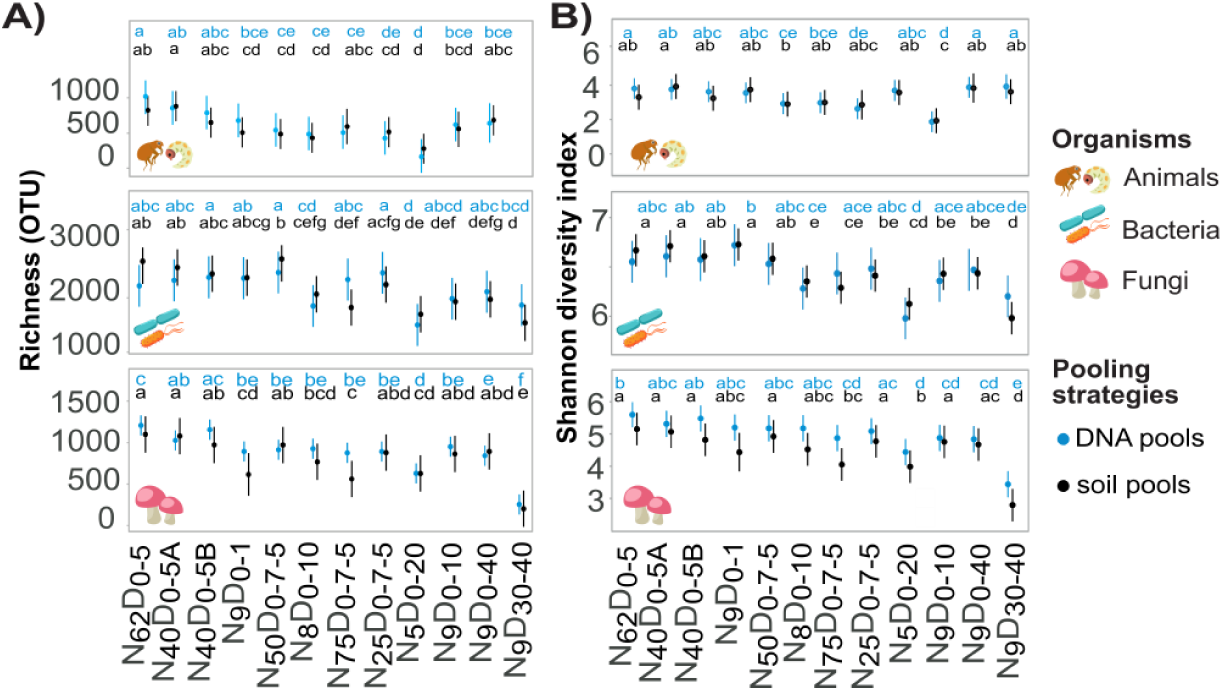
Within-site diversity captured by sample pooling designs. (A) Marginal means for OTU richness captured with soil and DNA pools across sampling designs in animal, bacterial and fungal communities. (B) Marginal means for Shannon diversity index. Linear models were used, with sampling sites and sampling designs treated as fixed factors. Similar letters mark sampling designs with statistically (α = 0.05) similar mean values.

In contrast to unpooled sampling, increasing the pooled subsamples from 40 to 62 did not significantly increase OTU richness. The lowest OTU richness in both DNA and soil pools was revealed with N_9_D30-40 and N_5_D_0-20_ designs. N_50_D_0-7.5_, N_25_D_0-7.5_ and N_75_D_0-7.5_ designs generally captured fewer OTUs than N_40_D_0-5_ designs in animals (up to 1.8-fold) and fungi (up to 6-fold). Meanwhile, the N_50_D_0-7.5_ design revealed slightly higher bacterial richness than other designs. Across all designs, the DNA pooling strategy revealed more animal and fungal OTUs than the soil pooling strategy (MC_DNA-SOIL_ = 0.09, p = 0.04; MC_DNA-SOIL_= 0.14, p = 0.02; see Table S5), but there was no effect in bacteria (MC_DNA-SOIL_= −0.08, p = 0.18).

### 3.4 Effect of sample pooling across different sequencing depths

To understand the pooling effect (PE) on diversity estimates across various sampling designs, we rarefied the pooled samples at the total sequencing depth of unpooled samples within the same design (i.e., 100% summary sequencing depth; Table S1) and defined PE as the ratio of OTU number between pooled and unpooled samples. It shows the gain (PE > 1) or loss (PE < 1) in OTU numbers captured with pooled compared to unpooled sampling. The range of PE_100%_ was 0.4–5.8, 0.5–2.4 and 0.6–1.7 in animals, bacteria and fungi, respectively, depending on sampling design and pooling strategy (Fig. 5A, B; Table S7). For the densely sampled N_62_D_0-5_, N_40_D_0-5A_ and N_40_D_0-5B_ designs, sample pooling caused OTU loss (PE_100%_ < 1) in animals and fungi. In contrast, DNA and only N_9_D30-40 soil pooling design enhanced fungal OTU recovery in other sampling designs (PE_100%_ > 1) in fungal communities. Soil and DNA pooling enhanced bacterial OTU recovery in all sampling designs, except for N_62_D_0-5_. Additionally, DNA and soil pooling only improved the ability of N5D_0-20_ and N_9_D_0-10_ to capture animal OTUs.

**Figure 5.**
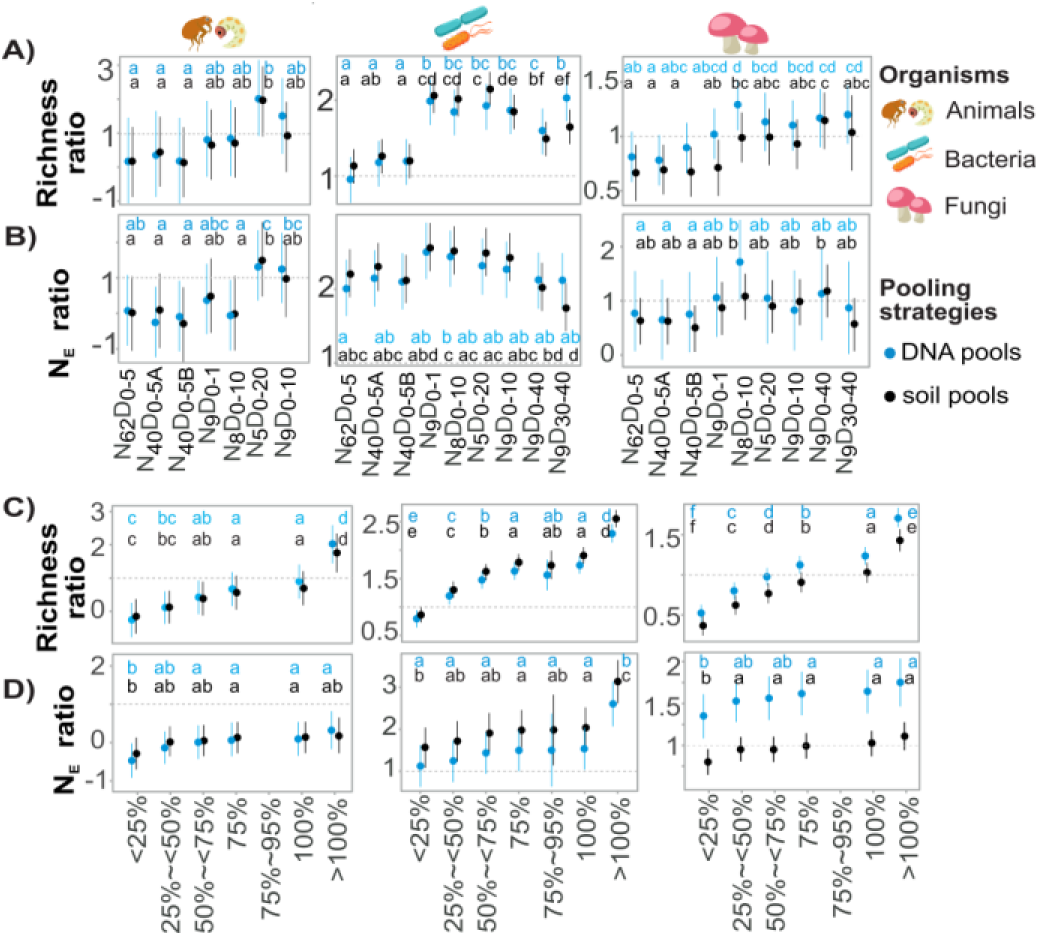
Pooling effect across sequencing depths and sampling designs. Pooling effect (ratio of within-site diversity estimates between pooled and unpooled samples) for (A) OTU richness and (B) effective number of OTUs across different sampling designs at the 100% summary sequencing depth. Pooling effect on (C) richness estimates and (D) effective number of OTUs (N_E_) across sequencing depths. The marginal means were derived from linear models, where sampling sites and sampling designs are treated as fixed factors. The x-axis represents the percentage of reads in pooled samples from the summary number of reads in unpooled samples within the same sampling design. Similar letters denote sampling designs with statistically (α = 0.05) similar mean values.

The PE varied with sequencing depth (Fig. 5C, D; see Table S7), differing among groups of organisms. The DNA and soil pooling did not improve animal diversity detection (PE < 1) at ≤ 100% summary sequencing depth. However, respectively, for bacteria and fungi, roughly 25% and 75% sequencing effort is enough to outperform unpooled sampling in OTU recovery.

The PE on richness was relatively higher in DNA pools than in soil pools for animals and fungi (animals: MC_SOIL-DNA_ = −0.2, p = 0.03; fungi: MC_SOIL-DNA_ = −0.08, p < 0.01; see Table S5), while soil pooling revealed relatively more bacterial OTUs (MC_SOIL-DNA_ = 0.08, p = 0.006). Based on the PE calculated for Shannon diversity indices, soil pooling generally retrieved more low-abundance fungal and bacterial OTUs than DNA pools, but there were no differences in animals.

Based on analyses performed for this study, we calculated that for the N_40_D_0-5_ design, sample pooling at the sampling stage (i.e., soil pooling) for 100 plots (4000 individual samples) saves 97.3% of resources destined for unpooled samples for chemicals, 76.9% of cost for sequencing library preparation plus 365 person-hours (97.3%) in the analysis of all animals, fungi and bacteria at the similar sequencing depth. For N_9_D_0-10_ and N_5_D_0-20_ designs, savings for labour are 74.7 (87.9%) and 37.4 (78.4%) person-hours, respectively, and savings for chemicals are 88.9% and 80%, respectively. Savings for sequencing are the same.

### 3.5 Contribution of molecular analysis artefacts in pooling effect

To understand the mechanisms of sample pooling effect on biodiversity estimates, we assessed whether PE could be attributable to rare species or undiscovered molecular analysis artefacts. We compared unpooled and pooled samples in the numbers and proportions of unique (detected in a site by only one design), rare (≤ 0.05% total reads) and artefactual OTUs at 100% summary sequencing depth. DNA pools, especially those with PE < 1, detected significantly fewer unique (up to 4.9 times) and rare (up to 1.47 times) OTUs and produced fewer unique artefacts (up to 11 times) compared to unpooled samples (Table 2 and S8). These patterns were especially pronounced in the Vasula sampling site. Soil pools did not differ in the numbers of rare, unique and artefactual OTUs from unpooled samples. Meanwhile, both DNA and soil pooling captured a smaller number of rare artefacts (up to 2 and 1.67 times for DNA and soil pools, respectively) but resulted in a higher proportion of rare artefacts (up to 1.6 times).

**Table 2.** Marginal contrasts (MC) in the proportion and counts of fungal rare, unique and artefactual OTUs between unpooled samples and DNA or soil pools.

| Parameters | Unpooled-DNA |  |  |  |  | Unpooled-soil |  |  |  |  |
| --- | --- | --- | --- | --- | --- | --- | --- | --- | --- | --- |
| | MC | p | Z value | R<sup>2</sup> | $\eta p^2$ | MC | p | Z value | R<sup>2</sup> | $\eta p^2$ |
| unique OTUs (PE > 1) | 5.1 | 0.07 | 1.8 | 0.33 | 0.1 | -0.2 | 0.9 | -0.1 | 0.4 | 0.001 |
| unique OTUs (PE < 1) | **18.4** | **< 0.001** | **6.4** | **0.8** | **0.7** | 6.5 | 0.07 | 1.8 | 0.4 | 0.1 |
| unique OTUs | **9.5** | **< 0.001** | **2.5** | **0.5** | **0.3** | 4.2 | 0.1 | 1.6 | 0.5 | 0.1 |
| rare OTUs (PE > 1) | 0.6 | 0.7 | 0.5 | 0.7 | 0.01 | 0.4 | 0.8 | 0.3 | 0.7 | 0.01 |
| rare OTUs (PE < 1) | **9.8** | **0.01** | **2.5** | **0.9** | **0.2** | 1.9 | 0.6 | 0.5 | 0.9 | 0.2 |
| rare OTUs/all OTUs | 0.01 | 0.8 | 0.3 | 0.4 | 0.01 | 0.02 | 0.4 | 0.9 | 0.4 | 0.01 |
| all artefacts/all OTUs | 0.02 | 0.4 | 0.9 | 0.5 | 0.03 | -0.01 | 0.6 | -0.5 | 0.5 | 0.03 |
| unique artefacts | **3.1** | **< 0.01** | **3** | **0.5** | **0.2** | 0.5 | 0.7 | 0.4 | 0.5 | 0.004 |
| rare artefacts | 1.8 | 0.3 | 0.3 | 0.7 | 0.02 | 1.4 | 0.4 | 0.8 | 0.7 | 0.02 |
| rare artefacts/rare OTUs | **-0.2** | **< 0.001** | **-3.6** | **0.3** | **0.2** | **-0.1** | **0.01** | **-2.5** | **0.3** | **0.2** |

## 4. Discussion

### 4.1 Sampling designs differ in diversity estimates

We showed that with unpooled sampling, the differences between sampling designs in richness estimates may exceed 27-fold. This depends on the taxonomic group of organisms and properties of the sampling design, including soil depth, number of subsamples and sampling area. Our analyses show that species of animals, bacteria and fungi continue accumulating with an increasing sampling effort of more than 200 individual samples, indicating that sampling effort contributes substantially to the differences in recovered diversity among sampling designs. In agreement, the sampling designs N_50_D_0-7.5_ and N_75_D_0-7.5_ recovered the highest number of species. Our extrapolation shows that to reach half the richness of a deeply sampled community (200 individual samples per site), 56–62, 48–53 and 43–49 samples would be needed for animals, bacteria and fungi from our study sites. We recommend at least 20 individual samples (see also Tedersoo et al. 2022), because designs with <10 samples were inconsistent in ranking the sites by richness.

Typically, the number of individual samples and sampling area are correlated (Dickie et al., 2018). Here, we specifically tested the importance of sampling area (i.e., plot size) on richness recovery based on subsampling the N_62_D_0-5_ design. Within the range of 327–2000 m^2^, the size of the sampling area did not affect the overall richness of soil biota. Similarly, reducing the sampling area from 2500 m² to 1400 m² using 40 subsamples resulted in comparable richness estimates. However, our layout of individual samples does not allow evaluation of whether the low diversity revealed with the N_5_D_0-20_ design is primarily due to an insufficient number of subsamples (N = 5) or to its exceptionally small area (16 m^2^) or the dilution effect of mineral soil below 5–10 cm. We recommend selecting an area large enough to enable collecting a sufficient number of samples (at least 20) outside the spatial autocorrelation range, i.e. a minimum area of 347 m^2^, 428 m^2^ and 645 m^2^ for 8 animal, bacterial and fungal individual samples, respectively.

Unlike Beule et al. (2022) and Eilers et al. (2012), we found that sampling within the top 5 cm or 10 cm is preferable for characterising the greatest part of the local soil biota. More deeply sampled soil columns harboured fewer species of fungi and bacteria compared with shallower ones. Subsoil samples harboured lower richness of all organism groups, corresponding to the previously observed vertical stratification of microbial diversity (Allison et al., 2007; Fierer et al., 2003; Queiroz et al., 2015; Schlatter et al., 2018). Furthermore, subsoil samples harboured a low proportion of unique OTUs, suggesting impoverished diversity, especially for fungi and animals. Because of the potential dilution effect (Du et al., 2017), adding mineral soil below 5 or 10 cm to molecular analysis may produce lower richness estimates. Nonetheless, OTU composition in relatively deep and shallow samples differs (Blume et al., 2002; Schlatter et al., 2018), but whether this can be mainly ascribed to the nestedness or turnover effects (Baselga, 2010; Martiny et al., 2011; Schlatter et al., 2018) remains unclear. The significant sampling depth effect indicates that the depth of sampling is a confounding parameter when comparing communities revealed using different sampling designs. Importantly, deeper sampling may reveal physicochemical properties that do not correspond to the sampled biota due to higher bulk density and lower microbial biomass of mineral soil.

### 4.2 Benefits and limitations of sample pooling

The effect of sample pooling on richness estimate accuracy differs among studies, depending on the sampling effort, ecosystem type, pooling strategy and taxonomic resolution of molecular methods (Adamo et al., 2021; Bullington et al., 2021; Engel et al., 2012; Grundmann & Gourbière, 1999; Manter et al., 2010; Ranjard et al., 2003). In our research, the pooling effect (the ratio between the pooled sample and individual samples in the captured site diversity) also differs among sampling designs, organisms of interest and sequencing depth. Specifically, the richness of fungi and especially animals was reduced by pooling, consistent with the results of Engel et al. (2012) and Manter et al. (2010). Meanwhile, in contrast to Manter et al. (2010) that found locally abundant bacterial species becoming undetectable in sample pools due to dilution, the richness of bacteria increased with pooling in our work.

We revealed that the pooling effect depends on the sampling design, particularly on the number of composed subsamples. Mixtures composed of >20 subsamples resulted in a loss of fungal and animal (but not bacterial) richness compared to unpooled sampling (i.e. PE < 1), irrespective of sequencing depth. Conversely, pooling <20 subsamples typically showed a gain in richness (PE > 1). As each individual sample is expected to include unique biological species because of their patchy distribution (O’Dwyer & Green, 2010; Ranjard & Richaume, 2001), we suggest that the loss of such unique and/or rare OTUs by sample pooling may contribute to the observed pooling effect. However, an in-depth analysis revealed a lower number of unique artefacts in DNA-pooled samples (but not soil-pooled samples) than in individual samples taken together. Because each amplicon may yield specific artefactual OTUs (Acinas et al., 2005), the sum of multiple individual amplicons likely contains a larger number of unique artefacts. It is possible that increasing chimera formation in diverse pooled samples (Omelina et al., 2019) is outweighed by summing up of artefacts in individual samples. Previous studies (Dopheide et al., 2019; Epp et al., 2012; Vesty et al., 2017) could not specifically address the remaining artefacts.

Although both DNA and soil pooling retained the imprint of sample size and soil depth on richness estimates, pooling resulted in an increase in bacterial and fungal community similarity among sampling designs. In addition, the differences among sampling designs in OTU richness or the effective number of OTUs were approximately 3-fold smaller compared with unpooled sampling. The amplification process may favour certain species and disfavour others through PCR bias (Acinas et al., 2005). Via a sampling effect, certain species in pooled samples are preferably amplified and sequenced, which may artificially homogenise communities in pooled samples.

The main criticism of pooling is the loss of information about within-site beta diversity and its local-scale drivers (Manter et al., 2010; Osborne et al., 2011). Using only pooled samples, we could not have estimated, for example, the spatial autocorrelation range, sampling area effect and optimal sample numbers. Our results also point to the homogenising effects of pooling, suggesting that biological differences among sites and ecological treatments may be more easily captured when using unpooled samples and combining sequencing data at a later stage. However, using pooled samples is feasible when comparing the data against other studies with different sampling designs due to the homogenisation effect.

The main advantage of sample pooling is the reduction of labour and analytical costs, particularly when assessing diversity across a large number of study sites (Baker et al., 2009; Osborne et al., 2011). For this study, the soil pooling saved up to 98.3% of resources, reducing labour and chemical costs 4.9-fold to 60.1-fold. Pooling affected DNA sequencing costs much less, because pooled samples may require deeper sequencing to reach PE > 1 (i.e., comprehensive sequencing depth). In spite of minor increases in richness, DNA pooling does not offer any advantage over soil pooling, because similar homogenising effects are evident, and DNA pooling has enormous additional costs of surplus DNA extraction. Indirectly, our analyses showed that lumping up to 62 DNA extracts of 0.2 grams of soil does not improve richness recovery for animals, bacteria or fungi when estimated based on a single DNA extraction from 0.2 grams of homogenised soil mixture. We admit that these inferences can be restricted to the sampled sites and encourage further data accumulation in other biomes using the sampling designs involved in this study as they base global- and continental-scale soil biodiversity projects (Tedersoo et al., 2021; Davison et al., 2012; Pärtel et al., 2019; Orgiazzi et al., 2018; Guerra et al., 2021; Delgado-Baquerizo et al., 2019; Maestre et al., 2012) that underlie the largest datasets (Delgado-Baquerizo et al., 2018; Mikryukov et al., 2023; Van Nuland et al., 2025) that are used in numerous studies on fungal and bacterial diversity and biogeography.

## Conclusions

For molecular biodiversity surveys of soil, representative sampling is necessary. Given the accumulation of soil biomass and biodiversity in the top few centimetres, sampling at a depth of 5 cm is sufficient to capture a large proportion of all animal, bacterial and fungal diversity. We recommend collecting at least 30 subsamples for animals, bacteria and fungi. Additional sampling effort depends on the intended sampling area, ease of vegetation penetration and heterogeneity of studied systems. The size of the sampling area had no effect on richness estimates at 327–2500 m^2^, but it should consider spatial autocorrelation range, potential edge effects and type of vegetation (i.e., trees vs. herbs). Considering labour and analytical costs, sample pooling in situ is feasible unless local-scale beta diversity or species co-occurrence analyses are intended. Considering the small but significant drawbacks of pooling on diversity, 2–3 smaller soil pools (<10 samples each) could be analysed separately, and the results summarised if the budget permits. Using pooled samples is also feasible when comparing the data against other studies with different sampling designs due to the homogenising effects.

## Acknowledgements

We thank Teresita M. Porter for her kind help in applying MetaWorks for COI data analysis. We thank Sten Anslan for his help in animal data filtering, Ali Hakimzadeh for supporting the fungal taxonomy in code and Liis Tiirmann for assistance in sampling. We thank Heidi Tamm for providing a breakdown of sequencing spending. This work was funded by the Estonian Science Foundation (grant PRG632) and the China Scholarship Council.

## Author Contributions

M.C. performed the analyses, visualized and interpreted the results and wrote the manuscript. V.M. provided analytical tools; O.D. visualized the results. O.C. provided animal data; L.T. designed the research, performed sampling and provided bacterial and fungal data. M.M., V.M., O.D. and M.C. built the interface. V.M., O.D. and L.T. supervised data analysis, participated in manuscript writing and results interpretation. All authors reviewed and edited the manuscript.

## Benefit-Sharing Statement

There are no benefits to report.

## Data Availability Statement

Demultiplexed Illumina and PacBio reads for each sample were deposited in the European Nucleotide Archive (ENA) under the accession number PRJEB88999. The calculator for sample processing costs is available at: https://mycology-microbiology-center.github.io/Soil_biodiversity_assessments/. All scripts used in the statistical analysis are available under the MIT license at GitHub: https://github.com/Mycology-Microbiology-Center/Soil_biodiversity_assessments.

## References

Acinas, S. G., Sarma-Rupavtarm, R., Klepac-Ceraj, V., & Polz, M. F. (2005). PCR-Induced Sequence Artifacts and Bias: Insights from Comparison of Two 16S rRNA Clone Libraries Constructed from the Same Sample. Applied and Environmental Microbiology, 71(12), 8966–8969. 10.1128/AEM.71.12.8966-8969.2005

Adamo, I., Piñuela, Y., Bonet, J., Castaño, C., Martínez de Aragón, J., Parladé, J., Pera, J., & Alday, J. (2021). Sampling forest soils to describe fungal diversity and composition. Which is the optimal sampling size in Mediterranean pure and mixed pine oak forests? Fungal Biology, 125. 10.1016/j.funbio.2021.01.005

Allison, V. J., Yermakov, Z., Miller, R. M., Jastrow, J. D., & Matamala, R. (2007). Using landscape and depth gradients to decouple the impact of correlated environmental variables on soil microbial community composition. Soil Biology and Biochemistry, 39(2), 505–516. 10.1016/j.soilbio.2006.08.021

Anderson, M. J. (2017). Permutational Multivariate Analysis of Variance (PERMANOVA). In Wiley StatsRef: Statistics Reference Online (pp. 1–15). 10.1002/9781118445112.stat07841

Anslan, S., Mikryukov, V., Armolaitis, K., Ankuda, J., Lazdina, D., Makovskis, K., Vesterdal, L., Schmidt, I. K., & Tedersoo, L. (2021). Highly comparable metabarcoding results from MGI-Tech and Illumina sequencing platforms. PeerJ, 9, e12254. 10.7717/peerj.12254

Arel-Bundock, V., Greifer, N., & Heiss, A. (2024). How to Interpret Statistical Models Using marginaleffects for R and Python. Journal of Statistical Software, 111(9), 1–32. 10.18637/jss.v111.i09

Bahram, M., Hildebrand, F., Forslund, S. K., Anderson, J. L., Soudzilovskaia, N. A., Bodegom, P. M., Bengtsson-Palme, J., Anslan, S., Coelho, L. P., Harend, H., Huerta-Cepas, J., Medema, M. H., Maltz, M. R., Mundra, S., Olsson, P. A., Pent, M., Põlme, S., Sunagawa, S., Ryberg, M., … Bork, P. (2018). Structure and function of the global topsoil microbiome. Nature, 560(7717), 233–237. 10.1038/s41586-018-0386-6

Baker, K. L., Langenheder, S., Nicol, G. W., Ricketts, D., Killham, K., Campbell, C. D., & Prosser, J. I. (2009). Environmental and spatial characterisation of bacterial community composition in soil to inform sampling strategies. Soil Biology and Biochemistry, 41(11), 2292–2298. 10.1016/j.soilbio.2009.08.010

Baselga, A. (2010). Partitioning the turnover and nestedness components of beta diversity. Global Ecology and Biogeography, 19(1), 134–143. 10.1111/j.1466-8238.2009.00490.x

Bates, D., Mächler, M., Bolker, B., & Walker, S. (2015). Fitting Linear Mixed-Effects Models Using lme4. Journal of Statistical Software, 67(1), 1–48. 10.18637/jss.v067.i01

Bell, T., Ager, D., Song, J.-I., Newman, J. A., Thompson, I. P., Lilley, A. K., & Gast, C. J. van der. (2005). Larger Islands House More Bacterial Taxa. Science, 308(5730), 1884–1884. 10.1126/science.1111318

Ben-Shachar, M. S., Lüdecke, D., & Makowski, D. (2020). effectsize: Estimation of Effect Size Indices and Standardized Parameters. Journal of Open Source Software, 5(56), 2815. 10.21105/joss.02815

Bengtsson-Palme, J., Ryberg, M., Hartmann, M., Branco, S., Wang, Z., Godhe, A., De Wit, P., Sánchez-García, M., Ebersberger, I., de Sousa, F., Amend, A., Jumpponen, A., Unterseher, M., Kristiansson, E., Abarenkov, K., Bertrand, Y. J. K., Sanli, K., Eriksson, K. M., Vik, U., … Nilsson, R. H. (2013). Improved software detection and extraction of ITS1 and ITS2 from ribosomal ITS sequences of fungi and other eukaryotes for analysis of environmental sequencing data. Methods in Ecology and Evolution, 4(10), 914–919. 10.1111/2041-210X.12073

Berg, M. P. (2012). 136Patterns of Biodiversity at Fine and Small Spatial Scales. In D. H. Wall, R. D. Bardgett, V. Behan-Pelletier, J. E. Herrick, T. H. Jones, K. Ritz, J. Six, D. R. Strong, & W. H. van der Putten (Eds.), Soil Ecology and Ecosystem Services (p. 0). Oxford University Press. 10.1093/acprof:oso/9780199575923.003.0014

Beule, L., Guerra, V., Lehtsaar, E., & Vaupel, A. (2022). Digging deeper: Microbial communities in subsoil are strongly promoted by trees in temperate agroforestry systems. Plant and Soil, 480(1), 423–437. 10.1007/s11104-022-05591-2

Blume, E., Bischoff, M., Reichert, J. M., Moorman, T., Konopka, A., & Turco, R. F. (2002). Surface and subsurface microbial biomass, community structure and metabolic activity as a function of soil depth and season. Applied Soil Ecology, 20(3), 171–181. 10.1016/S0929-1393(02)00025-2

Bokulich, N. A., Kaehler, B. D., Rideout, J. R., Dillon, M., Bolyen, E., Knight, R., Huttley, G. A., & Gregory Caporaso, J. (2018). Optimizing taxonomic classification of marker-gene amplicon sequences with QIIME 2’s q2-feature-classifier plugin. Microbiome, 6(1), 90. 10.1186/s40168-018-0470-z

Bolyen, E., Rideout, J. R., Dillon, M. R., Bokulich, N. A., Abnet, C. C., Al-Ghalith, G. A., Alexander, H., Alm, E. J., Arumugam, M., Asnicar, F., Bai, Y., Bisanz, J. E., Bittinger, K., Brejnrod, A., Brislawn, C. J., Brown, C. T., Callahan, B. J., Caraballo-Rodríguez, A. M., Chase, J., … Caporaso, J. G. (2019). Reproducible, interactive, scalable and extensible microbiome data science using QIIME 2. Nature Biotechnology, 37(8), 852–857. 10.1038/s41587-019-0209-9

Bradshaw, M. J., Aime, M. C., Rokas, A., Maust, A., Moparthi, S., Jellings, K., Pane, A. M., Hendricks, D., Pandey, B., Li, Y., & Pfister, D. H. (2023). Extensive intragenomic variation in the internal transcribed spacer region of fungi. iScience, 26(8). 10.1016/j.isci.2023.107317

Bruckner, A., Barth, G., & Scheibengraf, M. (2000). Composite sampling enhances the confidence of soil microarthropod abundance and species richness estimates. Pedobiologia, 44(1), 63–74. 10.1078/S0031-4056(04)70028-1

Bullington, L. S., Lekberg, Y., & Larkin, B. G. (2021). Insufficient sampling constrains our characterization of plant microbiomes. Scientific Reports, 11(1), 3645. 10.1038/s41598-021-83153-9

Callahan, B. J., McMurdie, P. J., Rosen, M. J., Han, A. W., Johnson, A. J. A., & Holmes, S. P. (2016). DADA2: High-resolution sample inference from Illumina amplicon data. Nature Methods, 13(7), 581–583. 10.1038/nmeth.3869

Chao, A., Henderson, P. A., Chiu, C.-H., Moyes, F., Hu, K.-H., Dornelas, M., & Magurran, A. E. (2021). Measuring temporal change in alpha diversity: A framework integrating taxonomic, phylogenetic and functional diversity and the iNEXT.3D standardization. Methods in Ecology and Evolution, 12(10), 1926–1940. 10.1111/2041-210X.13682

Chase, J. M., & Knight, T. M. (2013). Scale-dependent effect sizes of ecological drivers on biodiversity: Why standardised sampling is not enough. Ecology Letters, 16(s1), 17–26. 10.1111/ele.12112

Chiu, C.-H., & Chao, A. (2016). Estimating and comparing microbial diversity in the presence of sequencing errors. PeerJ, 4, e1634.

Davison, J., Öpik, M., Zobel, M., Vasar, M., Metsis, M., & Moora, M. (2012). Communities of Arbuscular Mycorrhizal Fungi Detected in Forest Soil Are Spatially Heterogeneous but Do Not Vary throughout the Growing Season. PLOS ONE, 7(8), e41938. 10.1371/journal.pone.0041938

Delgado-Baquerizo, M., Bardgett, R. D., Vitousek, P. M., Maestre, F. T., Williams, M. A., Eldridge, D. J., Lambers, H., Neuhauser, S., Gallardo, A., García-Velázquez, L., Sala, O. E., Abades, S. R., Alfaro, F. D., Berhe, A. A., Bowker, M. A., Currier, C. M., Cutler, N. A., Hart, S. C., Hayes, P. E., … Fierer, N. (2019). Changes in belowground biodiversity during ecosystem development. Proceedings of the National Academy of Sciences, 116(14), 6891–6896. 10.1073/pnas.1818400116

Delgado-Baquerizo, M., Oliverio, A. M., Brewer, T. E., Benavent-González, A., Eldridge, D. J., Bardgett, R. D., Maestre, F. T., Singh, B. K., & Fierer, N. (2018). A global atlas of the dominant bacteria found in soil. Science, 359(6373), 320–325. 10.1126/science.aap9516

Dickie, I. A., Boyer, S., Buckley, H. L., Duncan, R. P., Gardner, P. P., Hogg, I. D., Holdaway, R. J., Lear, G., Makiola, A., Morales, S. E., Powell, J. R., & Weaver, L. (2018). Towards robust and repeatable sampling methods in eDNA-based studies. Molecular Ecology Resources, 18(5), 940–952. 10.1111/1755-0998.12907

Dopheide, A., Xie, D., Buckley, T. R., Drummond, A. J., & Newcomb, R. D. (2019). Impacts of DNA extraction and PCR on DNA metabarcoding estimates of soil biodiversity. Methods in Ecology and Evolution, 10(1), 120–133. 10.1111/2041-210X.13086

Du, C., Geng, Z., Wang, Q., Zhang, T., He, W., Hou, L., & Wang, Y. (2017). Variations in bacterial and fungal communities through soil depth profiles in a Betula albosinensis forest. Journal of Microbiology, 55(9), 684–693. 10.1007/s12275-017-6466-8

Edgar, R. C., Haas, B. J., Clemente, J. C., Quince, C., & Knight, R. (2011). UCHIME improves sensitivity and speed of chimera detection. Bioinformatics, 27(16), 2194–2200. 10.1093/bioinformatics/btr381

Eilers, K. G., Debenport, S., Anderson, S., & Fierer, N. (2012). Digging deeper to find unique microbial communities: The strong effect of depth on the structure of bacterial and archaeal communities in soil. Soil Biology and Biochemistry, 50, 58–65. 10.1016/j.soilbio.2012.03.011

Engel, M., Behnke, A., Bauerfeld, S., Bauer, C., Buschbaum, C., Volkenborn, N., & Stoeck, T. (2012). Sample pooling obscures diversity patterns in intertidal ciliate community composition and structure. FEMS Microbiology Ecology, 79(3), 741–750. 10.1111/j.1574-6941.2011.01255.x

Epp, L. S., Boessenkool, S., Bellemain, E. P., Haile, J., Esposito, A., Riaz, T., Erséus, C., Gusarov, V. I., Edwards, M. E., Johnsen, A., Stenøien, H. K., Hassel, K., Kauserud, H., Yoccoz, N. G., Bråthen, K. A., Willerslev, E., Taberlet, P., Coissac, E., & Brochmann, C. (2012). New environmental metabarcodes for analysing soil DNA: potential for studying past and present ecosystems. Molecular Ecology, 21(8), 1821–1833. 10.1111/j.1365-294X.2012.05537.x

Fierer, N., & Jackson, R. B. (2006). The diversity and biogeography of soil bacterial communities. Proceedings of the National Academy of Sciences, 103(3), 626–631. 10.1073/pnas.0507535103

Fierer, N., Schimel, J. P., & Holden, P. A. (2003). Variations in microbial community composition through two soil depth profiles. Soil Biology and Biochemistry, 35(1), 167–176. 10.1016/S0038-0717(02)00251-1

Frøslev, T. G., Kjøller, R., Bruun, H. H., Ejrnæs, R., Brunbjerg, A. K., Pietroni, C., & Hansen, A. J. (2017). Algorithm for post-clustering curation of DNA amplicon data yields reliable biodiversity estimates. Nature Communications, 8(1), 1188. 10.1038/s41467-017-01312-x

Fukami, T. (2010). Community assembly dynamics in space. 10.1093/acprof:oso/9780199228973.003.0005

Green, J. L., Holmes, A. J., Westoby, M., Oliver, I., Briscoe, D., Dangerfield, M., Gillings, M., & Beattie, A. J. (2004). Spatial scaling of microbial eukaryote diversity. Nature, 432(7018), 747–750. 10.1038/nature03034

Green, R. H. (1979). Sampling design and statistical methods for environmental biologists. https://api.semanticscholar.org/CorpusID:63504719

Grundmann, L. G., & Gourbière, F. (1999). A micro-sampling approach to improve the inventory of bacterial diversity in soil. Applied Soil Ecology, 13(2), 123–126. 10.1016/S0929-1393(99)00027-X

Guerra, C. A., Bardgett, R. D., Caon, L., Crowther, T. W., Delgado-Baquerizo, M., Montanarella, L., Navarro, L. M., Orgiazzi, A., Singh, B. K., Tedersoo, L., Vargas-Rojas, R., Briones, M. J. I., Buscot, F., Cameron, E. K., Cesarz, S., Chatzinotas, A., Cowan, D. A., Djukic, I., van den Hoogen, J., … Eisenhauer, N. (2021). Tracking, targeting, and conserving soil biodiversity. Science, 371(6526), 239–241. 10.1126/science.abd7926

Jost, L. (2006). Entropy and diversity. Oikos, 113(2), 363–375. 10.1111/j.2006.0030-1299.14714.x

Kaehler, B. D., Bokulich, N. A., McDonald, D., Knight, R., Caporaso, J. G., & Huttley, G. A. (2019). Species abundance information improves sequence taxonomy classification accuracy. Nature Communications, 10(1), 4643. 10.1038/s41467-019-12669-6

Kang, S., & Mills, A. L. (2006). The effect of sample size in studies of soil microbial community structure. Journal of Microbiological Methods, 66(2), 242–250. 10.1016/j.mimet.2005.11.013

Katoh, K., & Standley, D. M. (2013). MAFFT Multiple Sequence Alignment Software Version 7: Improvements in Performance and Usability. Molecular Biology and Evolution, 30(4), 772–780. 10.1093/molbev/mst010

Kortmann, M., Chao, A., Schaefer, H. M., Blüthgen, N., Gelis, R., Tremlett, C. J., Busse, A., Püls, M., Seibold, S., Kriegel, P., Rabl, D., de la Hoz, M., Şekercioğlu, Ç. H., Schleuning, M., Feldhaar, H., Newell, F. L., Kümmet, S., Mitesser, O., Peters, M. K., & Müller, J. (2025). Sample coverage affects diversity measures of bird communities along a natural recovery gradient of abandoned agriculture in tropical lowland forests. Journal of Applied Ecology, 62(3), 480–491. 10.1111/1365-2664.14879

Kuznetsova, A., Brockhoff, P. B., & Christensen, R. H. B. (2017). lmerTest Package: Tests in Linear Mixed Effects Models. Journal of Statistical Software, 82(13), 1–26. 10.18637/jss.v082.i13

Larsson, A. (2014). AliView: A fast and lightweight alignment viewer and editor for large datasets. Bioinformatics, 30(22), 3276–3278. 10.1093/bioinformatics/btu531

Lemoine, N. P., Adams, B. J., Diaz, M., Dragone, N. B., Franco, A. L. C., Fierer, N., Lyons, W. B., Hogg, I. D., & Wall, D. H. (2023). Strong Dispersal Limitation of Microbial Communities at Shackleton Glacier, Antarctica. mSystems, 8(1), e01254–22. 10.1128/msystems.01254-22

Liu, G., Li, T., Zhu, X., Zhang, X., & Wang, J. (2023). An independent evaluation in a CRC patient cohort of microbiome 16S rRNA sequence analysis methods: OTU clustering, DADA2, and Deblur. Frontiers in Microbiology, Volume 14-2023. 10.3389/fmicb.2023.1178744

Lücking, R., Aime, M. C., Robbertse, B., Miller, A. N., Aoki, T., Ariyawansa, H. A., Cardinali, G., Crous, P. W., Druzhinina, I. S., Geiser, D. M., Hawksworth, D. L., Hyde, K. D., Irinyi, L., Jeewon, R., Johnston, P. R., Kirk, P. M., Malosso, E., May, T. W., Meyer, W., … Schoch, C. L. (2021). Fungal taxonomy and sequence-based nomenclature. Nature Microbiology, 6(5), 540–548. 10.1038/s41564-021-00888-x

Lüdecke, D., Ben-Shachar, M. S., Patil, I., & Makowski, D. (2020). Extracting, Computing and Exploring the Parameters of Statistical Models using R. Journal of Open Source Software, 5(53), 2445. 10.21105/joss.02445

Lüdecke, D., Ben-Shachar, M. S., Patil, I., Waggoner, P., & Makowski, D. (2021). performance: An R Package for Assessment, Comparison and Testing of Statistical Models. Journal of Open Source Software, 6(60), 3139. 10.21105/joss.03139

MacArthur, R. H. (1965). Patterns of species diversity. Biological Reviews, 40(4), 510–533. 10.1111/j.1469-185X.1965.tb00815.x

Maestre, F. T., Quero, J. L., Gotelli, N. J., Escudero, A., Ochoa, V., Delgado-Baquerizo, M., García-Gómez, M., Bowker, M. A., Soliveres, S., Escolar, C., García-Palacios, P., Berdugo, M., Valencia, E., Gozalo, B., Gallardo, A., Aguilera, L., Arredondo, T., Blones, J., Boeken, B., … Zaady, E. (2012). Plant Species Richness and Ecosystem Multifunctionality in Global Drylands. Science, 335(6065), 214–218. 10.1126/science.1215442

Manter, D. K., Weir, T. L., & Vivanco, J. M. (2010). Negative Effects of Sample Pooling on PCR-Based Estimates of Soil Microbial Richness and Community Structure. Applied and Environmental Microbiology, 76(7), 2086–2090. 10.1128/AEM.03017-09

Martiny, J. B. H., Eisen, J. A., Penn, K., Allison, S. D., & Horner-Devine, M. C. (2011). Drivers of bacterial β-diversity depend on spatial scale. Proceedings of the National Academy of Sciences, 108(19), 7850–7854. 10.1073/pnas.1016308108

McMurdie, P. J., & Holmes, S. (2013). phyloseq: An R Package for Reproducible Interactive Analysis and Graphics of Microbiome Census Data. PLOS ONE, 8(4), 1–11. 10.1371/journal.pone.0061217

Mikryukov, V. (2025). metagMisc: Miscellaneous functions for metagenomic analysis. 10.5281/zenodo.597622

Mikryukov, V., Anslan, S., & Tedersoo, L. (2025). NextITS: a pipeline for metabarcoding eukaryotes with full-length ITS sequenced with PacBio (Version 1.0.0) [Computer software]. 10.5281/zenodo.15074881

Mikryukov, V., Dulya, O., Zizka, A., Bahram, M., Hagh-Doust, N., Anslan, S., Prylutskyi, O., Delgado-Baquerizo, M., Maestre, F. T., Nilsson, H., Pärn, J., Öpik, M., Moora, M., Zobel, M., Espenberg, M., Mander, Ü., Khalid, A. N., Corrales, A., Agan, A., … Tedersoo, L. (2023). Connecting the multiple dimensions of global soil fungal diversity. Science Advances, 9(48), eadj8016. 10.1126/sciadv.adj8016

Minh, B. Q., Schmidt, H. A., Chernomor, O., Schrempf, D., Woodhams, M. D., von Haeseler, A., & Lanfear, R. (2020). IQ-TREE 2: New Models and Efficient Methods for Phylogenetic Inference in the Genomic Era. Molecular Biology and Evolution, 37(5), 1530–1534. 10.1093/molbev/msaa015

Morita, H., & Akao, S. (2021). The effect of soil sample size, for practical DNA extraction, on soil microbial diversity in different taxonomic ranks. PLOS ONE, 16(11), 1–15. 10.1371/journal.pone.0260121

Nilsson, R. H., Tedersoo, L., Ryberg, M., Kristiansson, E., Hartmann, M., Unterseher, M., Porter, T. M., Bengtsson-Palme, J., Walker, D. M., de Sousa, F., Gamper, H. A., Larsson, E., Larsson, K.-H., Kõljalg, U., Edgar, R. C., & Abarenkov, K. (2015). A Comprehensive, Automatically Updated Fungal ITS Sequence Dataset for Reference-Based Chimera Control in Environmental Sequencing Efforts. Microbes and Environments, 30(2), 145–150. 10.1264/jsme2.ME14121

Oksanen, J., Blanchet, F. G., Kindt, R., Legendre, P., Minchin, P., O’Hara, B., Simpson, G., Solymos, P., Stevens, H., & Wagner, H. (2015). Vegan: Community Ecology Package. R Package Version 2.2-1, 2, 1–2

Omelina, E. S., Ivankin, A. V., Letiagina, A. E., & Pindyurin, A. V. (2019). Optimized PCR conditions minimizing the formation of chimeric DNA molecules from MPRA plasmid libraries. BMC Genomics, 20(7), 536. 10.1186/s12864-019-5847-2

Orgiazzi, A., Ballabio, C., Panagos, P., Jones, A., & Fernández-Ugalde, O. (2018). LUCAS Soil, the largest expandable soil dataset for Europe: A review. European Journal of Soil Science, 69(1), 140–153. 10.1111/ejss.12499

Osborne, C. A., Zwart, A. B., Broadhurst, L. M., Young, A. G., & Richardson, A. E. (2011). The influence of sampling strategies and spatial variation on the detected soil bacterial communities under three different land-use types. FEMS Microbiology Ecology, 78(1), 70–79. 10.1111/j.1574-6941.2011.01105.x

O’Dwyer, J. P., & Green, J. L. (2010). Field theory for biogeography: A spatially explicit model for predicting patterns of biodiversity. Ecology Letters, 13(1), 87–95. 10.1111/j.1461-0248.2009.01404.x

Pärtel, M., Carmona, C. P., Zobel, M., Moora, M., Riibak, K., & Tamme, R. (2019). DarkDivNet – A global research collaboration to explore the dark diversity of plant communities. Journal of Vegetation Science, 30(5), 1039–1043. 10.1111/jvs.12798

Porter, T. M., & Hajibabaei, M. (2022). MetaWorks: A flexible, scalable bioinformatic pipeline for high-throughput multi-marker biodiversity assessments. PLOS ONE, 17(9), 1–11. 10.1371/journal.pone.0274260

Porter, T. M., Morris, D. M., Basiliko, N., Hajibabaei, M., Doucet, D., Bowman, S., Emilson, E. J. S., Emilson, C. E., Chartrand, D., Wainio-Keizer, K., Séguin, A., & Venier, L. (2019). Variations in terrestrial arthropod DNA metabarcoding methods recovers robust beta diversity but variable richness and site indicators. Scientific Reports, 9(1), 18218. 10.1038/s41598-019-54532-0

Quast, C., Pruesse, E., Yilmaz, P., Gerken, J., Schweer, T., Yarza, P., Peplies, J., & Glöckner, F. O. (2013). The SILVA ribosomal RNA gene database project: Improved data processing and web-based tools. Nucleic Acids Research, 41(D1), D590–D596. 10.1093/nar/gks1219

Queiroz, C., Silva, F., & Rossa-Feres, D. (2015). The relationship between pond habitat depth and functional tadpole diversity in an agricultural landscape. Royal Society Open Science, 2. 10.1098/rsos.150165

Querner, P., & Bruckner, A. (2010). Combining pitfall traps and soil samples to collect Collembola for site scale biodiversity assessments. Applied Soil Ecology, 45(3), 293–297. 10.1016/j.apsoil.2010.05.005

R Core Team. (2023). R: A Language and Environment for Statistical Computing. R Foundation for Statistical Computing. https://www.R-project.org/

Rahbek, C. (2005). The role of spatial scale and the perception of large-scale species-richness patterns. Ecology Letters, 8(2), 224–239. 10.1111/j.1461-0248.2004.00701.x

Ranjard, L., Lejon, D. P. H., Mougel, C., Schehrer, L., Merdinoglu, D., & Chaussod, R. (2003). Sampling strategy in molecular microbial ecology: Influence of soil sample size on DNA fingerprinting analysis of fungal and bacterial communities. Environmental Microbiology, 5(11), 1111–1120. 10.1046/j.1462-2920.2003.00521.x

Ranjard, L., & Richaume, A. (2001). Quantitative and qualitative microscale distribution of bacteria in soil. Research in Microbiology, 152(8), 707–716. 10.1016/S0923-2508(01)01251-7

Robeson, M. S., II, O’Rourke, D. R., Kaehler, B. D., Ziemski, M., Dillon, M. R., Foster, J. T., & Bokulich, N. A. (2021). RESCRIPt: Reproducible sequence taxonomy reference database management. PLOS Computational Biology, 17(11), e1009581. 10.1371/journal.pcbi.1009581

Rognes, T., Flouri, T., Nichols, B., Quince, C., & Mahé, F. (2016). VSEARCH: a versatile open source tool for metagenomics. PeerJ Preprints, 4, e2409v1. 10.7287/peerj.preprints.2409v1

Schlatter, D. C., Kahl, K., Carlson, B., Huggins, D. R., & Paulitz, T. (2018). Fungal community composition and diversity vary with soil depth and landscape position in a no-till wheat-based cropping system. FEMS Microbiology Ecology, 94(7), fiy098. 10.1093/femsec/fiy098

Song, Z., Schlatter, D., Kennedy, P., Kinkel, L. L., Kistler, H. C., Nguyen, N., & Bates, S. T. (2015). Effort versus Reward: Preparing Samples for Fungal Community Characterization in High-Throughput Sequencing Surveys of Soils. PLOS ONE, 10(5), 1–13. 10.1371/journal.pone.0127234

Tan, L., Fan, C., Zhang, C., von Gadow, K., & Fan, X. (2017). How beta diversity and the underlying causes vary with sampling scales in the Changbai mountain forests. Ecology and Evolution, 7(23), 10116–10123. 10.1002/ece3.3493

Tedersoo, L., Bahram, M., Zinger, L., Nilsson, R.H, Kennedy, P., Yang, T., Anslan, S., Mikryukov, V. (2022). Best practices in metabarcoding of fungi: from experimental design to results. Molecular Ecology, 31(12), 2769–2795

Tedersoo, L., Hosseyni Moghaddam, M. S., Mikryukov, V., Hakimzadeh, A., Bahram, M., Nilsson, R. H., Yatsiuk, I., Geisen, S., Schwelm, A., Piwosz, K., Prous, M., Sildever, S., Chmolowska, D., Rueckert, S., Skaloud, P., Laas, P., Tines, M., Jung, J.-H., Choi, J. H., … Anslan, S. (2024). EUKARYOME: the rRNA gene reference database for identification of all eukaryotes. Database, 2024, baae043. 10.1093/database/baae043

Tedersoo, L., Mikryukov, V., Anslan, S., Bahram, M., Khalid, A. N., Corrales, A., Agan, A., Vasco-Palacios, A.-M., Saitta, A., Antonelli, A., Rinaldi, A. C., Verbeken, A., Sulistyo, B. P., Tamgnoue, B., Furneaux, B., Ritter, C. D., Nyamukondiwa, C., Sharp, C., Marín, C., … Abarenkov, K. (2021). The Global Soil Mycobiome consortium dataset for boosting fungal diversity research. Fungal Diversity, 111(1), 573–588. 10.1007/s13225-021-00493-7

Van Nuland, M. E., Averill, C., Stewart, J. D., Prylutskyi, O., Corrales, A., van Galen, L. G., Manley, B. F., Qin, C., Lauber, T., Mikryukov, V., Dulia, O., Furci, G., Marín, C., Sheldrake, M., Weedon, J. T., Peay, K. G., Cornwallis, C. K., Větrovský, T., Kohout, P., … SPUN Mapping Consortium. (2025). Global hotspots of mycorrhizal fungal richness are poorly protected. Nature, 645(8080), 414–422. 10.1038/s41586-025-09277-4

Vesty, A., Biswas, K., Taylor, M. W., Gear, K., & Douglas, R. G. (2017). Evaluating the Impact of DNA Extraction Method on the Representation of Human Oral Bacterial and Fungal Communities. PLOS ONE, 12(1), e0169877. 10.1371/journal.pone.0169877

Wang, Q., Garrity George M., Tiedje James M., & Cole James R. (2007). Naïve Bayesian Classifier for Rapid Assignment of rRNA Sequences into the New Bacterial Taxonomy. Applied and Environmental Microbiology, 73(16), 5261–5267. 10.1128/AEM.00062-07

Whitcomb, S., & Stutz, J. C. (2007). Assessing diversity of arbuscular mycorrhizal fungi in a local community: Role of sampling effort and spatial heterogeneity. Mycorrhiza, 17(5), 429–437. 10.1007/s00572-007-0118-5

Zhang, H., & Bu, W. (2022). Exploring Large-Scale Patterns of Genetic Variation in the COI Gene among Insecta: Implications for DNA Barcoding and Threshold-Based Species Delimitation Studies. Insects, 13(5). 10.3390/insects13050425

