## Supplemental figures for "Sampling design and sample processing affect soil biodiversity assessments"

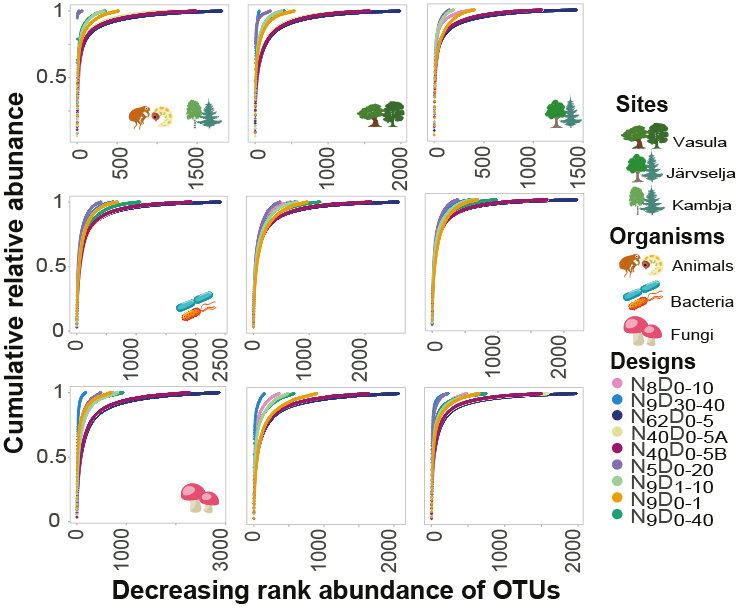

**Figure S1.** **Within-site diversity across unpooled sampling designs.** The relative cumulative abundance curves in animal, bacterial, and fungal communities. OTU numbers were ordered by decreasing relative abundance within each sampling design.

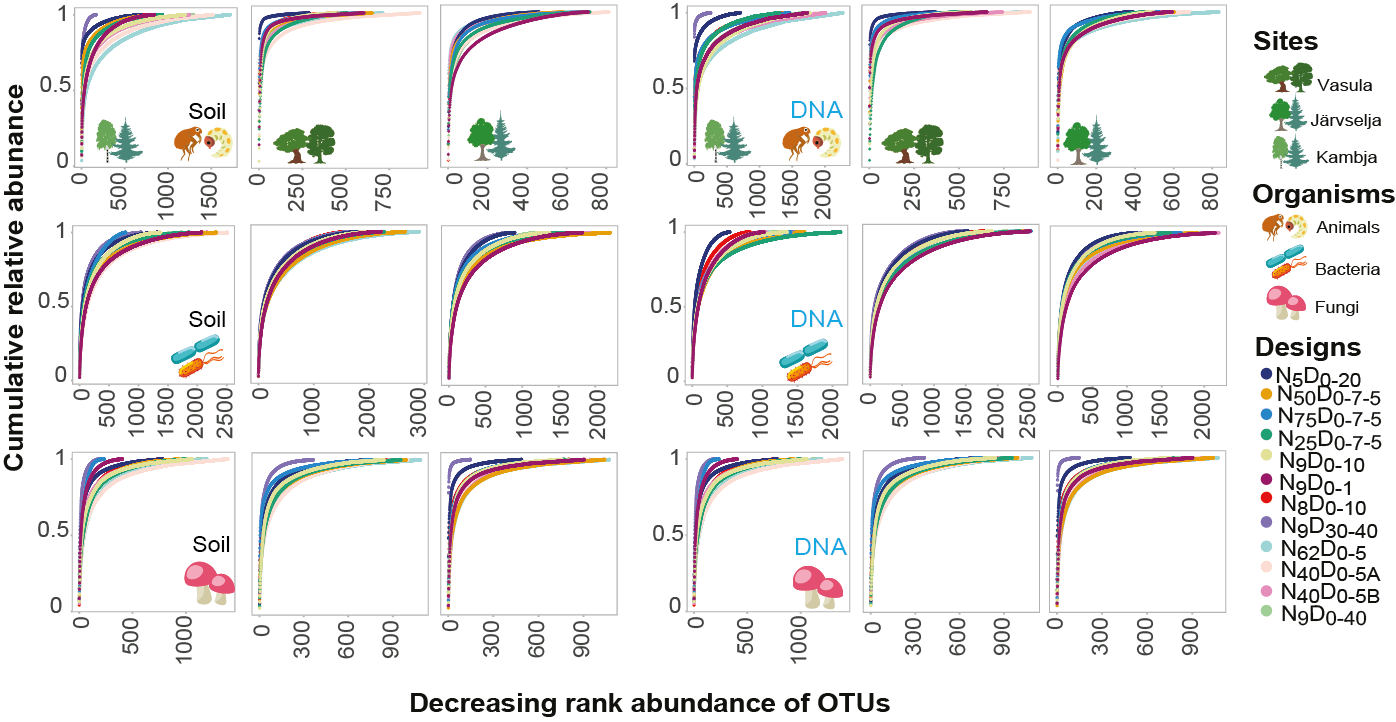

**Figure S2.** **Within-site diversity across pooling sampling designs.** The relative cumulative abundance curves in animal, bacterial, and fungal communities. OTU numbers were ordered by decreasing relative abundance within each sampling design.

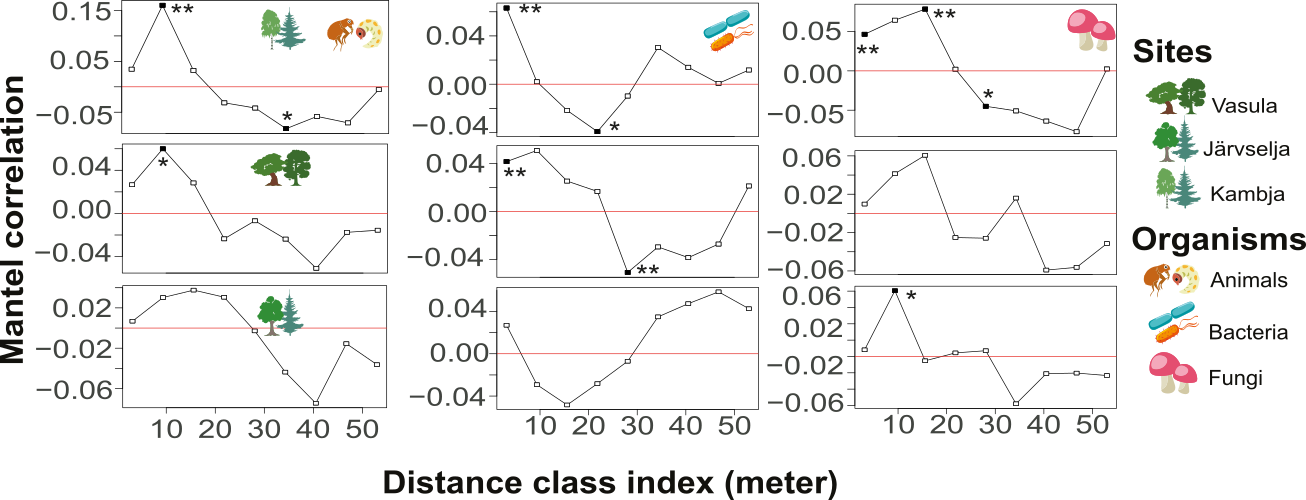

**Figure S3. Spatial Correlation in Communities.** This figure illustrates the spatial autocorrelation for animal, bacterial and fungal communities. Significant correlation values indicate that communities separated by corresponding distance (see X axis) show similarity (positive correlation values) or dissimilarity (negative correlation values). Asterisks highlight statistically significant correlations.
